# Bursts of regional cortical inhibition during smartphone use

**DOI:** 10.1101/2025.08.24.671850

**Authors:** Wenyu Wan, Arko Ghosh

**Affiliations:** Cognitive Psychology Unit, Institute of Psychology, Leiden University, The Netherlands; Department of Psychology, University of Amsterdam, The Netherlands

## Abstract

Transient burst of beta (β) oscillations – brief neural events lasting only a fraction of a second – are increasingly recognised for their role in functional inhibition that may support cognitive processes and motor timing underlying goal-directed behaviour. Smartphone behaviour, which involves sustained engagement across diverse cognitive processes provides a fresh avenue to examine these bursts. In this study, we used EEG to measure β-bursts as participants freely interacted with their smartphones using their right hand for ∼ 80 min. Bursts were detected across the scalp, with bilateral sensorimotor electrodes accumulating more periods with bursts than the rest of the electrodes. There was a brain-wide reduction in the probability of bursts preceding a touchscreen touch, followed by an increase, with the strongest modulation in the left sensorimotor electrodes. Interestingly, touchscreen touches sometimes occurred during the bursts. We further examined the intra-burst touch rate across the scalp and found that the rate was lower over the left sensorimotor cortex than the rest of electrodes. Finally, we characterised the diverse behaviours in terms of the interval between touchscreen interactions to find that the behaviours with bursts were typically longer than the behaviours without bursts. We propose that the apparently continuous stream of smartphone touches is supported by brief, spatially localised transient bursts of inhibition. These transients may help support the computational flexibility in distributed neural networks needed for processing continuous motor output and integrating the rich sensory information.

## Introduction

Rhythmic brain dynamics in the β-band (∼13–30 Hz) range is correlated to various cognitive processes from top-down sensory processing to the precise timing of motor outputs (Engel & Fries, 2010; Khanna & Carmena, 2015). There is an emerging idea that these oscillations – and activity in other frequency bands – transiently occur in bursts rather than being sustained over time (Lundqvist et al., 2024; van Ede et al., 2018). This perspective offers new clues on the functional role of β-band apart from offering a more accurate description of the physiological signals (Engel & Fries, 2010; Lundqvist et al., 2024). Both the conventional literature on β activity and the emerging burst measurements show a reduction in sensorimotor β activity surrounding movements followed by a rebound - although the bursts appear more spatially localised and are better predictors of behaviour than the conventional measures (Feingold et al., 2015; Khanna & Carmena, 2017; Little et al., 2019; Tzagarakis et al., 2010). As the bursts appear across the cortex one attractive idea is that they generally contribute to top-down neural processing by enabling transient and localised inhibition that supports goal-directed behaviour (Lundqvist et al., 2024). Moreover, the transients may enable inter-regional communication necessary for coordinating multiple cognitive processes and integrating information and controlling motor outputs (Little et al., 2019; Lundqvist et al., 2018). The rich behaviours expressed in the real-world recruit multiple cognitive processes and involve complex event sequences. Capturing β bursts in such naturalistic settings may be helpful to uncovering their role in coordinating distributed computational processes across large-scale neural networks.

Smartphone behaviour engages large and distributed neural populations. There are scattered efforts to leverage the quantifiable nature of the behaviour to address the underlying cognitive process (Balerna & Ghosh, 2018; Gindrat et al., 2015; Kock et al., 2023; van de Ruit & Ghosh, 2022; Wan et al., 2024). The fluctuations in the amount of touchscreen interactions from one minute to the next reflect in the β-band power such that higher the use lower the power over the sensorimotor electrodes (van de Ruit & Ghosh, 2022). Although the behaviour involves a complex train of touchscreen interactions, time-locking the β-band power fluctuations to the touch reveals suppression around smartphone interactions across the scalp but predominantly over the sensorimotor electrodes (Kock et al., 2023). It remains unclear, however, whether these seemingly gradual modulations in average β power reflect changes in the probability of β bursts, rather than sustained oscillatory activity.

The idea that β bursts offer brief and local inhibition to support top-down cognitive processes underlying goal-directed behaviour, raises a range of questions on their role in smartphone behaviour. First, smartphone touchscreen interactions are rather multi-processes involving sensory and motor processing. It is established that the location of the β bursts appear to be task-specific (Lundqvist et al., 2024). During smartphone behaviour, do the bursts span several cortical areas? Second, β bursts in the sensorimotor cortex are linked to motor inhibition (Wessel, 2020). Given that touchscreen interactions are uninstructed, how does β burst probability fluctuate around the touches – can spontaneous behavioural outputs emerge during ongoing somatosensory β bursts? Third, smartphone behaviour is continuous – with the user moving from one touchscreen interaction to the next with varying interval timing. For the idea that the bursts enable a “clear out” of working memory or recently evoked sensorimotor processes (Schmidt et al., 2019), does this process occur after each touchscreen touch? Alternatively, are the transients more likely to occur for certain smartphone behaviours over others – for instance, do the behaviours characterised by long touchscreen interaction intervals show more bursts than those with shorter intervals?

Addressing an array of questions rest on measuring and analysing β bursts during smartphone behaviour. Here, we captured the transient neural patterns in 54 individuals by using EEG as they used their own smartphones in the laboratory for a period of ∼ 80 minutes. During the EEG recording, the participants were simply instructed to conduct smartphone activities that they commonly performed in the real world. We captured the spatial distribution of the bursts by recording across the scalp. We analysed the bursts time-locked to the smartphone touchscreen events to reveal how the burst probability fluctuated surrounding the touch. Finally, we captured the nature of the smartphone behaviour – in terms of touchscreen interaction dynamics – in the presence of bursts. Our findings reveal a temporal and spatial distribution of β bursts aligned to the spontaneous smartphone behavioural dynamics.

## Results

### β-burst occupancy during smartphone use

We detected β-bursts while individuals interacted with their smartphone. The duration of the bursts varied across participants, electrodes and burst events. At the population level, median burst durations ranged from 149 to 178 ms across the scalp, based on per-electrode medians within participants. We estimated the β-burst occupancy (BO) – a proportion of recording time spent in bursts (*sum of burst durations / total recording time*).

The bursts were observed across the scalp ranging from an occupancy from 0.10 to 0.15 in the population level (**Fig. 1**). The bursts were common in the sampled population as indicated by the one-sample t-tests against 0 (where a null value indicates an absence of bursts, **Fig. 1**). To address the topology of the bursts we z-normalised the BOs across the scalp. This transformation followed by a one-sample t-test revealed a relatively high occupancy of burst over the bilateral sensorimotor cortex and a low occupancy at the posterior electrodes (**Fig. 1**).

**Figure 1.**
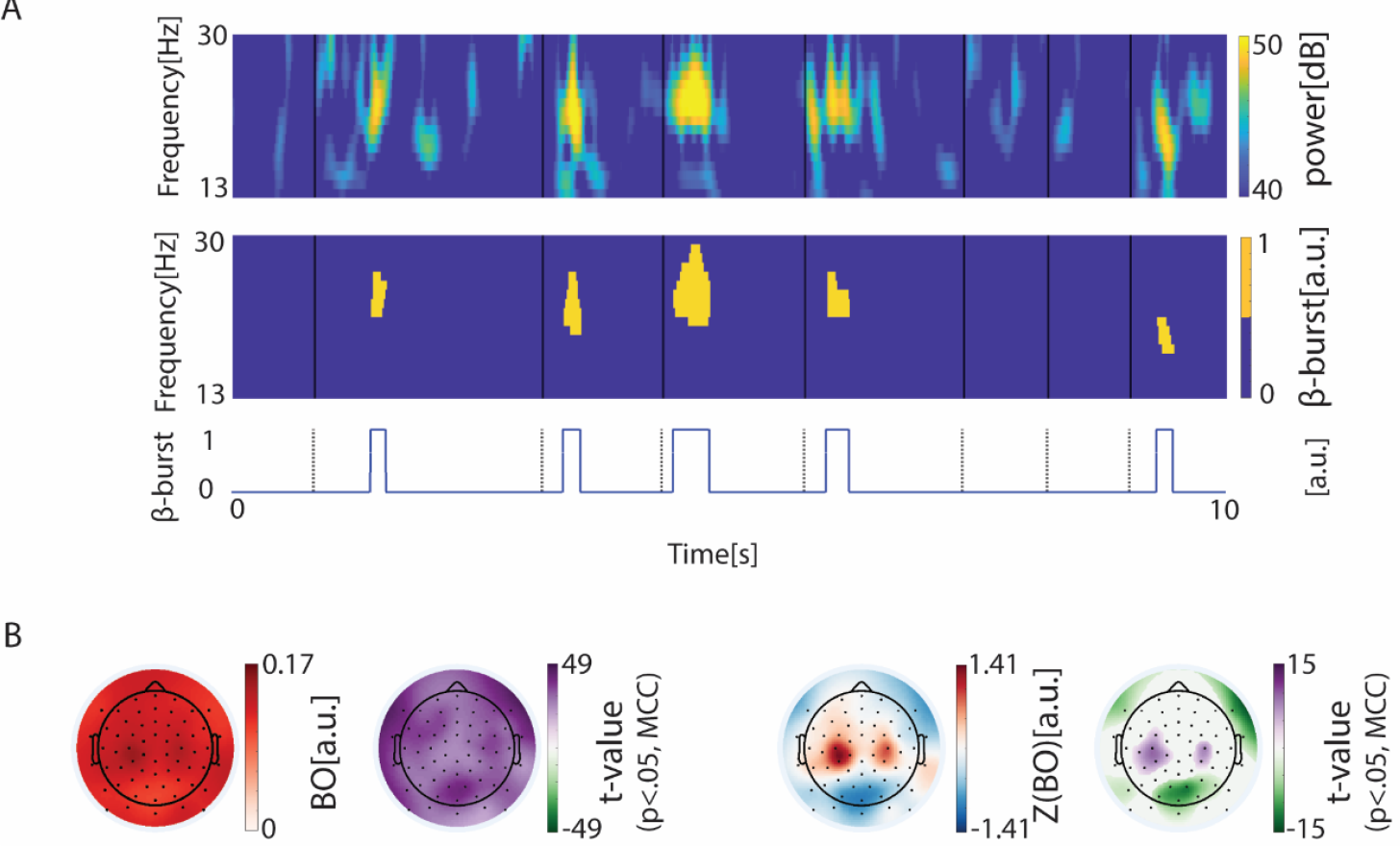
Spatial distribution of β-burst occupancy (in continuous EEG) during smartphone use. (A) β-burst detection from continuous time-frequency activity. The top row shows 10 seconds of continuous beta-band (13–30 Hz) time-frequency activity. The middle row displays the corresponding binary β-burst detection based on power and duration thresholds. The bottom row shows a simplified binary representation, indicating the occurrence of β-bursts. (B) The left panel exhibits the grand-averaged β-bursts occupancy (BO, calculated as the total burst duration divided by recording time per electrode) across the population, along with t-statistics from one-sample t-tests against zero, corrected for multiple comparisons using the Bonferroni method. The right panel has the same legend but displays results based on the z-normalized BO values.

### β-bursts time-locked to the smartphone touchscreen touch

Smartphone use consists of a complex sequence of touchscreen events (median inter-touch interval ∼871 ms, iqr ∼1236 ms). We next addressed if and how the β-bursts are aligned to the isolated touchscreen events. Towards this, we time-locked the bursts to the touchscreen event yielding a raster from which we determined the β-burst probability index (BPI) spanning -3 s to 3 s from the touch (**Fig. 2A**). This revealed a fluctuation of bursts in the immediate surroundings of the touch, such that the bursts were diminished ∼1685 ms prior to the touch to rebound by ∼992 ms after the touch for the somatosensory electrode across the population. A similar pre-touch suppression was also observed at frontal electrodes, albeit with different timing (**Fig. 2B**).

**Figure 2.**
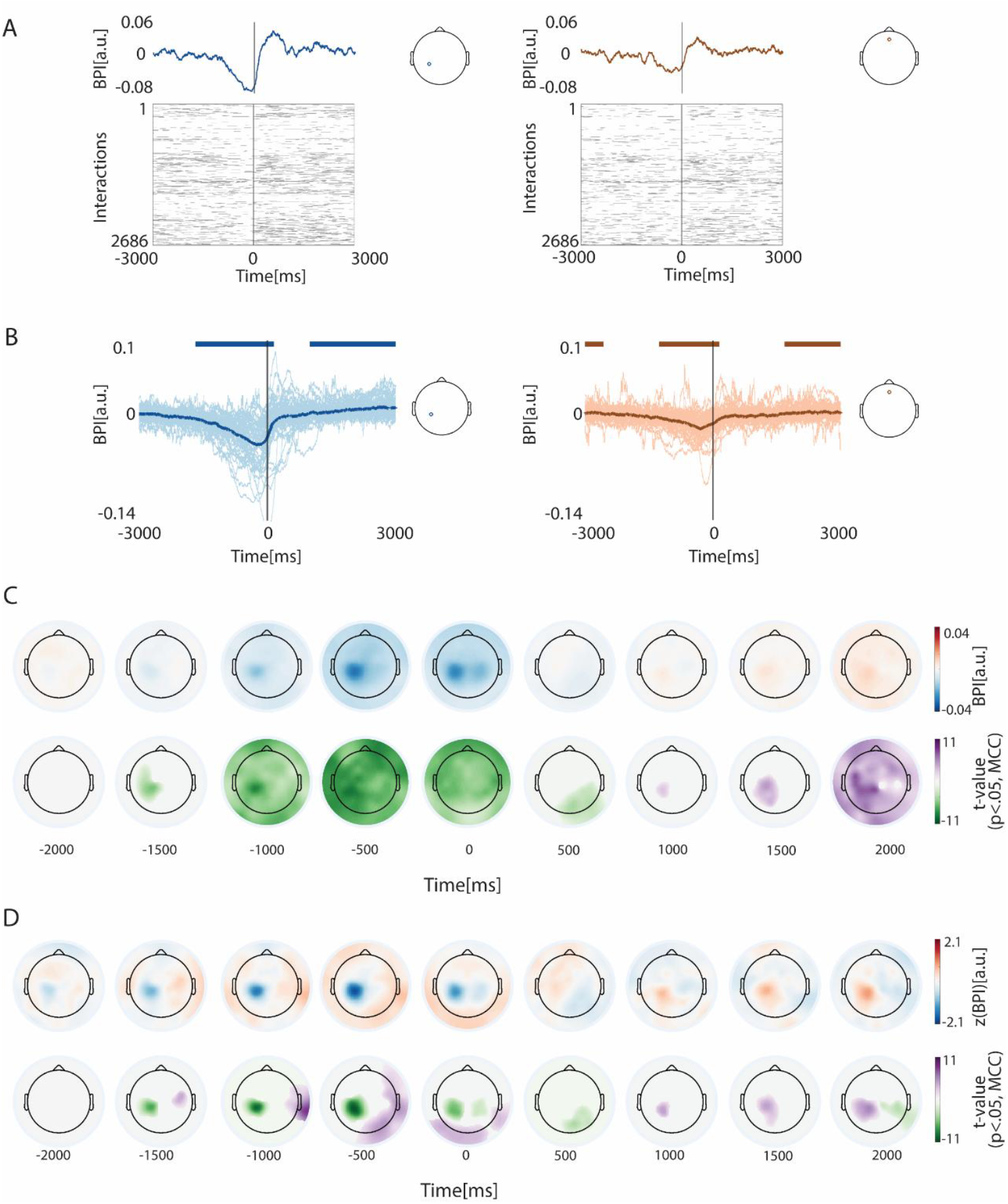
β-bursts time-locked to smartphone touchscreen events. (A) Top: Baseline corrected β-burst probability estimated from a raster of bursts time-locked to the touchscreen touch. Data from one representative participant and two electrodes (left: contralateral sensorimotor electrode, right: frontal electrode, location as indicated in inserts). Bottom: Trial-by-trial binarized β-burst activity from –3 to +3 s relative to touchscreen touch. (B) Bold lines show the grand-averaged time-locked β-burst probability across participants for two electrodes (left: frontal; right: left sensorimotor, as shown in inserts). Thin lines represent individual participants. Top horizontal bars indicate time points with baseline corrected β-burst probabilities significantly different from zero according to one-sample t-test (p<0.05, MCC). (C) Top row: scalp topography of grand-averaged time-locked β-burst probability index (BPI) across participants at selected time points relative to interaction onset. Bottom row: t-statistic from one-sample t-test results comparing BPI to zero, corrected for multiple comparison correction using spatial-temporal clustering based on 1000 bootstraps (α = 0.05). (D) Top row: same results with (C), but z-normalized across electrodes. Bottom row: same t-statistic results, but based on the z-normalized BPIs.

We addressed the time-locked fluctuations across the electrodes. This revealed a scalpwide reduction in bursts starting at ∼1100 ms prior to the touch to rebound by ∼2000 ms (**Fig. 2C**). We used z-transformations to reveal the differences between the electrodes. According to one-sample t-tests, the left sensorimotor electrodes (contralateral to the movement) showed relatively strong burst suppression prior to the touch whereas the same electrodes showed a rebound more than the rest of the electrodes after the touch. We quantified the fluctuations in the immediate surrounding of the touch (-1s to 1s) in terms of the extent and timing of the burst reduction and rebound (**Fig. 3**). The minimum burst probabilities were confined to the left (contralateral) somatosensory cortex. In terms of the relative time of this drop, the right somatosensory cortex along with the temporal electrode was the latest to reach the minimum. In terms of the maximum burst in this period, the somatosensory electrodes showed the lowest values but appeared the earliest compared to the other electrodes.

**Figure 3.**
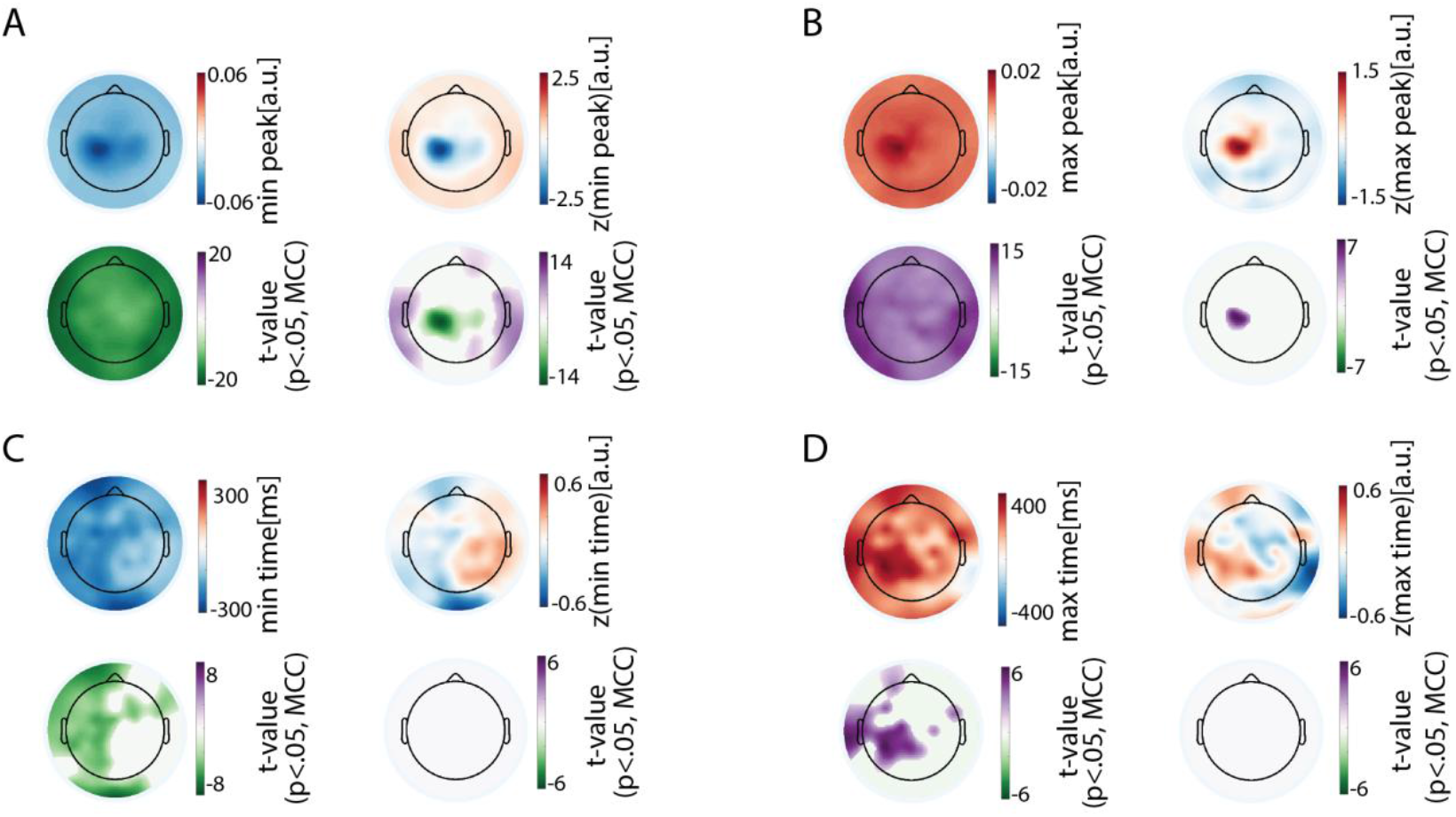
Magnitude and timing of β-burst suppression and rebound surrounding smartphone touchscreen events. (A) Left panel shows the spatial distribution of the minimum (suppression) β-burst probability across the population. The adjacent topography shows the corresponding t-statistics from one-sample t-tests against zero, masked at *p* <.05 with Bonferroni correction for multiple comparisons (MCC). The right panel shows the same results but based on z-normalized values across the scalp. (B) Same as in (A), but for the maximum (rebound) β-burst probability. (C) Left panel shows the timing of the minimum β-burst probability (suppression) and the corresponding t-statistics from one-sample t-tests, masked at *p* <.05 (MCC). Right panel shows the same results but based on the z-normalized timing values. (D) Same as in (C), but for the timing of the maximum β-burst probability (rebound).

In sum, there was considerable fluctuations in β-bursts surrounding isolated smartphone touchscreen events. While there was brain-wide modulation in bursts surrounding the touch, the fluctuations were relatively exaggerated over the contralateral somatosensory cortex – both in terms of burst suppression and a quick rebound.

### Smartphone touchscreen events surrounding β-bursts

The analysis of burst time-locked to smartphone touchscreen events revealed fluctuations in burst probability, with a reduction prior to the touch. But these probabilities before baseline correction never really dropped to zero, suggesting that burst may play a more flexible role in tuning behaviour outputs. To further examine this possibility, we performed a series of analysis to characterize how bursts accompany behavioural output.

First, we estimated the touch rate when the bursts were *ON* by dividing the number of touches occurring during β-burst by the total β-burst duration. While the behaviour did occur during bursts detected across the scalp, it was least probable for the bursts over the contralateral sensorimotor cortex and most probable for the burst over the occipital and temporal electrodes (**Fig. 4A**). Finally, we separated the smartphone touchscreen interaction intervals according to whether a burst occurred in the interval or not. We quantified the difference between the interval duration distributions as β-burst behavioural timing index (BBTI). In general, the intervals were longer in the presence of bursts than without across the whole scalp. When z-transformed, the contralateral sensorimotor electrodes showed relatively large gaps whereas the right parietal electrode revealed the smallest gaps (**Fig. 4C**).

**Figure 4.**
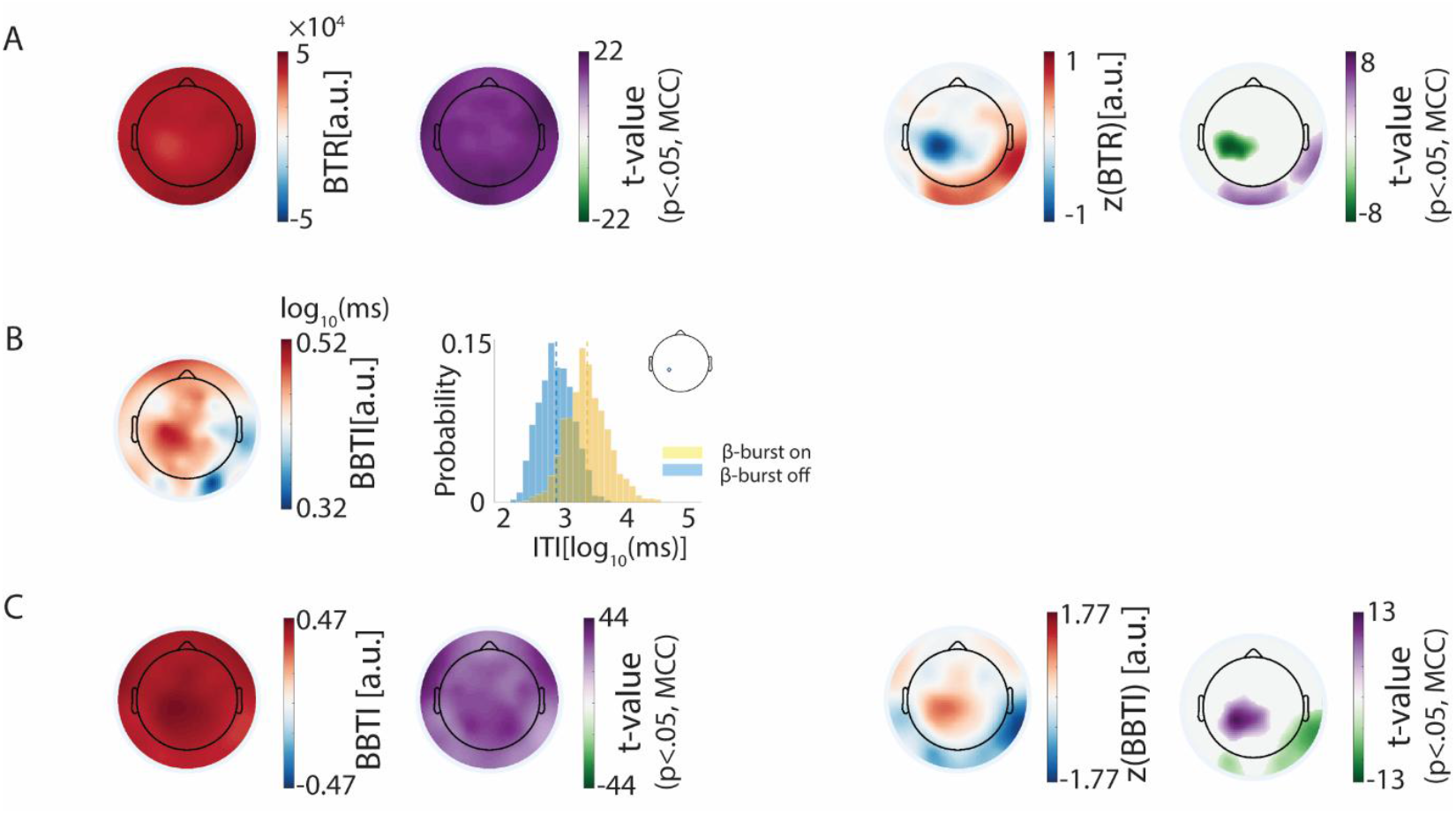
β-burst modulate the likelihood and timing of smartphone behaviour (A) Behaviour probability when β-burst across scalp. Left panel shows the topography of grand averaged interactions probability (z-transformed) during β-burst. Right panel shows the t-statistic result from one-sample t-test against zero (masked by p <.05, MCC). (B) Left panel shows the temporal pattern difference (log_10_ scale) of behaviour with and without β-burst across scalp from one participant (BBTI). Right panel shows the distribution of inter-interaction intervals with and without β-burst for the sensorimotor electrode (as insert shown). (C) Left panel shows the topography of grand-averaged intervals distribution difference between with and without β-burst (BBTI) across participants and the t-statistic of one-sample t-test against zero (masked by p <.05, MCC). Right panel shows the same results with left panel, but based on the z-normalized BBTIs.

In sum, these results consistently suggest that β-bursts do not strictly suppress movement but rather modulate both the likelihood and timing of behaviour output by dynamically shifting between on and off states in a region-specific manner.

## Discussion

We found β-bursts across the scalp when engaged in smartphone behaviour. Although smartphone behaviour involves a range of cognitive processes – including sensorimotor processing, vision and memory – there was a concentration of β-bursts and corresponding fluctuations in the sensorimotor cortex. Smartphone behaviour consists of a continuous train of touchscreen interactions. Although each touch is embedded in a train of behavioural outputs, we find a momentary reduction of burst probability surrounding an individual touch. Still, touches can be generated during the burst-ON periods and they were more likely to be observed in behaviours with longer inter-touch intervals. These patterns suggest that during real-world behaviour spatio-temporal distribution of transient inhibitory neural processes support the variety of parallel processes underlying smartphone behaviour.

In general, there was a pronounced concentration of β-bursts in the sensorimotor cortex and this was in keeping in prior observations where bilateral sensorimotor bursts are predominant may it be at rest or during behaviour (Little et al., 2019; Power & Bardouille, 2021; Rayson et al., 2023; Wessel, 2020). The distinct circuitry of the sensorimotor cortex and how it is driven by subcortical structures may partly explain the concentration of bursts (Sherman et al., 2016; Tinkhauser et al., 2018; Yu et al., 2021). Alternatively, it may reflect the unique nature of peripheral somatosensory afferents (Sherman et al., 2016; Tuthill & Azim, 2018). Here, the β-bursts serve as a gating or filtering mechanism (Liljefors et al., 2024), continuously regulating cortical access to afferent sensory input from the sensory thalamus (Baker, 1971; Karvat et al., 2021).

Surrounding the individual touchscreen touches, we found that the burst probability across the scalp diminished and then rebounded in keeping with what has been observed before using conventional measures of beta activity surrounding motor responses (Jurkiewicz et al., 2006; Kock et al., 2023; Little et al., 2019; Torrecillos et al., 2018). This supports the idea that the fluctuations in beta activity previously attributed to sustained changes occur in the form of transient changes in neural activity. One widely held view is that β-band activity is associated with maintaining the current cognitive or motor status (Engel & Fries, 2010). From this perspective, the β-burst probability reduction before touches may reflect a release from the existing status, allowing a shift in cognitive activity or behaviour. The β-burst probability increases after touches may reflect the reestablishment of stability. From one touch to the next, the timing of the bursts showed substantial variation, raising the possibility that β-bursts are timed according to a complex mixture of intrinsic – such as cortical excitability (Samaha et al., 2017) – and extrinsic factors – such as kinematics of the thumb (Kock et al., 2023).

Beta activity in the sensorimotor cortex has been implicated in motor inhibition – as if the inhibition must be overcome to enable a movement (Wessel, 2020). In our study, we found the modulation of burst probability surrounding touchscreen touches appeared to support this inhibitory role. In particular, the sensorimotor bursts were the least permissive to the touchscreen touches indicating that inhibition there may be the hardest to overcome for smartphone touch. However, we did observe touchscreen touches sometimes occurred during periods of β-bursts. One possibility is that, beyond motor inhibition and in parallel to it, β-bursts in the sensorimotor network also support functions such as sensory prediction, integration, and motor planning (Little et al., 2019; Shin et al., 2017; Tatz et al., 2023; West et al., 2023). The longer burst durations accompanying smartphone touches too may stem from a composite of these support functions. Another possibility is that β-bursts do not strictly serve as inhibitory gates to stop movement but rather act to slow down or delay behavioural output. Supporting this possibility, the smartphone intervals between touchscreen touches with bursts in the sensorimotor cortex were longer than the intervals without the bursts.

There may be an alternative explanation as to why the intervals were longer with the bursts than without. The behaviours with longer intervals may be supported by more diverse cognitive processes than in the shorter intervals. The bursts may be necessary to coordinate these diverse processes spanning well beyond the sensorimotor cortex (Betancourt et al., 2023; Hannah et al., 2020; Haufler et al., 2022; Kopell et al., 2000; Liljefors et al., 2024; Lundqvist et al., 2018; Sporn et al., 2020; Tatz et al., 2023). In our observations the gap between the interval distributions with vs. without burst spanned across the scalp albeit the gap was the most prominent in the sensorimotor cortex. This is consistent with the idea that the inhibitory transients reflected in the β-bursts are used across the scalp for coordinating neural processes (Lundqvist et al., 2024). Our observation helps link this idea to the diverse behaviours captured on the smartphone.

In conclusion, brain wide β-bursts accompany smartphone interactions. β-bursts are typically studied in laboratory designed tasks with a clear onset and offset. They are a signal of interest in clinical research focused on movement disorders and for instance, used to explain parkinsonian symptoms (Cagnan et al., 2019; Lofredi et al., 2019; Tinkhauser, Pogosyan, Little, et al., 2017; Tinkhauser, Pogosyan, Tan, et al., 2017; Tinkhauser et al., 2018; Vinding et al., 2020). The continuous behavioural outputs captured on the smartphone offer a fresh perspective on these inhibitory transients. In the ongoing behaviour β-bursts fluctuated surrounding a specific event – the touchscreen touch. Future research to manipulate the bursts can help clarify whether they causally influence the behaviour (Pogosyan et al., 2009). β-bursts has been implicated in a range of processes – from clearing WM to action inhibition. The inseparable mixture of executive and motor processes in continuous and spontaneous smartphone behaviour offers a new vertical to study β-bursts. Their distribution in a ubiquitous behaviour may not only help address basic questions on the shared neural mechanisms spanning different cortical areas but also help yield a new marker for abnormal brain functions.

## Method

### Participants

We recruited 64 participants from agestudy.nl research platform (Ceolini et al., 2022). This set of participants has been previously reported in Wan and Ghosh (2025). All participants reported that they had no history of neurological or psychiatric disorders. Four participants were excluded because of the updated self-reported unhealthy status soon after the measurement session. Two participants were excluded due to missing smartphone data during EEG measurement for unknown reasons. Four participants were excluded were excluded due to the poor clock alignment across the different instrumentation (see details below). All participants provided informed consent. This study was approved by the Ethics Committee of Psychology at Leiden University (ERB-reference number: 2020-02-14-Ghosh, dr. A.-V2-2044).

### Smartphone data collection

Smartphone behavior was recorded using the TapCounter app (QuantActions AG, Switzerland) (Balerna & Ghosh, 2018), which was installed on participant’s smartphone. The app continuously logged the timestamps of all smartphone touchscreen events with millisecond precision in UTC, along with the label of the app in use. Participants used their smartphone freely with the right thumb (as instructed) during the EEG measures conducted here for a period of ∼80 min.

### Movement sensor recording and clock alignment

Movement data were recorded using a movement sensor (Flex Sensor, 112 mm, Digi-Key, Thief River Falls). The sensor was attached to the dorsum of the participant’s right thumb. This setup allowed participants to interact with smartphone freely. The analogue signals from the sensor were sampled at 1 Hz using the Polybox (Brain Products GmbH, Gilching, Germany), and were stored in the EEG dataset file, sharing the same clock as the EEG recordings.

Movement signals were processed following Kock et al. (2023). In brief, the raw signals were filtered between 1 and 10 Hz. The continuous signal was then epoched ranging from -3 to 3 s relative to the onset of smartphone touchscreen touches. For each participant, movement signals were averaged across all touches.

Since the smartphone clock may differ from the laboratory clock, we corrected for clock differences following the procedure described in previous studies (Kock et al., 2023; Wan & Ghosh, 2025). Briefly, a pre-trained model was used to estimate the screen touch times based on the movement signal. Then, the time delay between the predicted touches and the smartphone-recorded touches was corrected. For participants who have more than one recording sessions (say induced by a bathroom break or impedance check), the predicted touch time may slightly differ across sessions because of kinematic differences between sessions. We then aligned the movement waveforms using *alignsignals* from Matlab’s Signal processing toolbox (MathWorks, Natick, MA, USA).

### EEG recording and preprocessing

EEG data was collected by using 64 channels actiCap Snap cap (62 scalp electrodes, 2 ocular electrodes) with customized equidistant layout (Brain Products GmbH, Gilching, Germany). The EEG signals were gathered referenced to the vertex and amplified with BrainAmp amplifier (Brain Products GmbH, Gilching, Germany). The sampling rate of recorded and digitalized signals was 1 kHz. Participants were asked to arrive for the measurement with washed hair and scalp. The skin at the contact sites were further degreased by using alcohol swabs. Supervisc gel (Easycap GmbH, Herrsching, Germany) was applied to obtain an electrical contact between the skin and electrode, and targeted an impedance under 10 kΩ for each electrode. The entire EEG measurement lasted approximately 1.5 hours in total.

We used the EEGLAB toolbox (Delorme & Makeig, 2004) and custom scripts (shared on GitHub) on MATLAB 2023b (MATLAB, Mathworks, Natick) to pre-process the EEG data. EEG data was high pass filtered at 0.5 Hz. The behaviourally inactive data segments – determined based on touchscreen touches – were removed if both pre-and post-touchscreen event intervals were longer than 30 s. Adaptive mixed independent component analysis (Palmer et al., 2012; Palmer et al., 2008) was used to decompose independent component. Based on AMICA results, we rejected artifact components with over than 90% likelihood originating from artifacts like eyeblinks, muscle, etc., as evaluated by ICLabel toolbox (Pion-Tonachini et al., 2019). EEG data was further band-pass filtered with 1-45 Hz. We rejected bad channels with the correlation less than 0.85 with neighbouring channels by *clean_channels*. Rejected channels were interpolated using the spherical interpolation method by *pop_interp*.

### β-burst detection

We used the β-burst detection procedure as described in (Shin et al., 2017; Wessel, 2020). Briefly, a complex Morlet wavelet approach was used to decompose the time-frequency (TF) power on the continuous EEG data spanning 13 to 30 Hz (Fig. 1A). An individual β-burst was detected when power was greater than x 6 the median of power spanning the whole recording period. β-burst durations less than 2 cycles for each frequency were further excluded. We then obtained a binary β-burst TF matrix. From this binary burst TF matrix, we further derived a binary β-burst time series.

### Parameters based on β-bursts

#### β-burst occupancy (BO)

We calculated the β-burst occupancy based on the continuous binary β-burst time series data, defined as the total burst duration divided by the whole recording duration. The calculation was performed for each participant across all electrodes. To study the differences between the electrodes, the β-burst occupancy was z-normalized across the electrodes. For population level statistics, we pooled either the BBO or the z-normalised values and subsequently conducted a one-sample t-test against zero using limo_eeg toolbox (Pernet et al., 2011), followed by Bonferroni correction for multiple comparisons.

#### Time locked β-burst probability index (BPI)

We calculated the BPI for each participant and electrode by epoching the continuous binary β-burst time series from -3 to 3 s relative to the onset of smartphone touch, averaging across trials, and correcting baseline by subtracting the median value across the time. BPIs were normalized across electrodes to emphasize the difference in spatial distribution for each time point. Population-level analyses were performed by pooling BPIs or the z-normalised values across participants, and conducting one-sample t-tests against zero, with multiple comparisons corrected via spatial-temporal clustering (1000 bootstraps, α = 0.05). Topographies suggested variability in the timing and extent of β-burst suppression across electrodes. To capture this variability across electrodes, we identified the minimum value of the BPI within the – 1 to 1 s window (relative to touch onset) for each electrode, along with the corresponding time point. We also normalized those values across the scalp to emphasize spatial variation. We pooled all raw and z-transformed values of suppression timing and magnitude across participants and performed one-sample t-tests against zero, with Bonferroni correction for multiple comparisons. A similar analysis was conducted to characterize β-burst rebound by identifying the maximum value and corresponding time point within the -1 to 1 s window relative to touch onset. One-sample t-tests were performed on the pooled raw and z-transformed values of rebound timing and magnitude, followed by Bonferroni correction.

#### Intra-burst touch rate (BTR)

From the time-locked β-burst probability index result, we observed that β-bursts sometimes occurred during periods of β-bursts. To quantify how extent β-bursts co-occur with touches, we calculated the burst-on touch rate by dividing the number of touches that occurred during β-bursts by the total β-burst duration. The BTRs were then z-transformed across the scalp to emphasize spatial difference. One-sample t-test against zero, followed by Bonferroni multiple comparison correction, was conducted on the pooled BTRs and z-transformed values.

#### β-burst behaviour timing index (BBTI)

To examine how β-burst relate to behavioural timing, we separated touchscreen intervals with and without the bursts and compared the timing of behaviour between these two conditions. In brief, we separated the binary β-burst time series based on inter-touch intervals. Each data segment was categorized as with β-burst group if any burst occurred within the interval or categorized as without β-burst group otherwise. We then calculated the difference in the median behaviour timing (log_10_ scale) between those two groups for each electrode as BBTI. The BBTIs were then normalized across electrodes. We performed one-sample t-tests against zero on the pooled BBTIs and z-transformed values across the population, followed by Bonferroni multiple comparison correction.

## Data, Materials, and Software Availability

The data will be available on dataverse.nl upon publication. Custom-written scripts for preprocessing analysis are available at: https://github.com/CODELABCODELIB/JID_ERP_Smartphone_2024, and scripts for beta burst identification and subsequent analyses are shared at: https://github.com/CODELABCODELIB/BetaburstInhibition_2025.

## Acknowledgements

This study was funded by Velux Stiftung (Grant No.1283, awarded to A.G. with Richard Ridderinkhof as co-applicant). We acknowledge the assistance of Beste Yavuz, Lysanne Groenewegen, Barbora Michalidesová, Lorenzo Van Hoorde in EEG data collection. We also grateful to all participants for their contribution to this study.

## Author contributions

A.G. conceived the study. A.G. and W.W. designed the study. W.W acquired, performed EEG preprocessing, and conducted the analyses with the aid of A.G. W.W drafted the manuscript with the aid of A.G.. A.G., W.W. edited the manuscript.

